# NetBoxR: Automated Discovery of Biological Process Modules by Network Analysis in R

**DOI:** 10.1101/2020.06.02.129387

**Authors:** Eric Minwei Liu, Augustin Luna, Guanlan Dong, Chris Sander

## Abstract

**Summary:** Large-scale sequencing projects, such as The Cancer Genome Atlas (TCGA) and the International Cancer Genome Consortium (ICGC), have accumulated a variety of high throughput sequencing and molecular profiling data, but it is still challenging to identify potentially causal genetic mutations in cancer as well as in other diseases in an automated fashion. We developed the NetBoxR package written in the R programming language, that makes use of the NetBox algorithm to identify candidate cancer-related processes. The algorithm makes use of a networkbased approach that combines prior knowledge with a network clustering algorithm, obviating the need for and the limitation of functionally curated gene sets. A key aspect of this approach is its ability to combine multiple data types, such as mutations and copy number alterations, leading to more reliable identification of functional modules. We make the tool available in the Bioconductor R ecosystem for applications in cancer research and cell biology.

**Availability and implementation:** The NetBoxR package is free and open-sourced under the GNU GPL-3 license R package available at https://www.bioconductor.org/packages/release/bioc/html/netboxr.html

**Supplementary information:** None

## Introduction

Large-scale sequencing consortia such as The Cancer Genome Atlas (TCGA) (Cancer Genome Atlas Research Network et al., 2013) and the Interactional Cancer Genome Consortium (ICGC) (International Cancer Genome Consortium et al., 2010) provide detailed genomic alteration profiling in many cancer types. Many methods based on the recurrence of genomic alterations, i.e., the frequency of occurrence in sets of tumor samples, have been developed to identify alterations likely to be functional in oncogenesis or cancer progression, addressing an important question in the field of precision oncology (Bailey et al., 2018). However, due to the considerable patient-to-patient heterogeneity of the cancer genome, rare mutations in certain patients can still be involved in tumor development by affecting biological processes in ways similar to those of known cancer genes. One way to address the issue of the effect of rare mutations is to combine prior knowledge of genetic and molecular interactions with recurrence-based methods and thus increase the power of predictions in spite of relatively low recurrence counts. In this spirit, we have developed the NetBox algorithm that seeks to automate the identification of candidate oncogenic processes and involved genes, which allows the quantitative analysis of genomic alterations in the context of known signaling pathway connectivity (Cerami et al., 2010). The NetBox algorithm identifies potentially novel network modules by mapping genomic alterations onto a comprehensive prior-knowledge interaction network, containing nodes and their interactions (edges), and then identifying modules as clusters of connected nodes that are frequently affected by genomic alterations as a set. The aggregation of nodes into clusters overcomes the statistical problem of low counts for individual nodes. This is in contrast to methods such as gene set enrichment analysis (GSEA), a popular approach to associate a gene list to biological functions, that relies on curated, pre-defined clusters of genes and does not make use of the often known interactions between genes or gene products. Unlike GSEA, NetBox is not limited to nor influenced by curated gene lists, it takes into account known interactions, and overcomes the issue of the occurrence of genes in more than one of the curated gene sets. Instead, the NetBox algorithm derives network modules de novo, based on the alteration data in tumor samples, such that the identified modules cannot only recover or overlap with known predefined pathways but, more importantly, identify new functional gene groups that cross curated gene boundaries. For newly discovered modules, the functional annotation can then be derived from the annotation of the gene members, facilitating discoveries.

To extend the use of NetBox, we have implemented the NetBox algorithm as a native R package, NetBoxR. The NetBoxR package provides users with access to the NetBox algorithm within the R ecosystem, thereby providing simplicity and flexibility for the visualization and secondary analyses through available R packages by using common data structures in R packages. Here, we describe the use of NetBoxR to integrate various types of genomic alterations for the detection of potentially functional network modules in glioblastoma multiforme (GBM) cancer, as an example, and highlight NetBoxR tutorial material for integrating the use of the NetBoxR package with additional R packages.

## Methods

### Implementation

The NetBoxR package implements the original NetBox algorithm for discovery of pathway modules (Cerami et al., 2010) using the R programming language and adds several functions to communicate with other packages in the R ecosystem and to integrate several input data types

### Base functionality and algorithm

NetBoxR takes significantly altered genes from SNV mutation, copy number alteration associated with gene expression change, and merges them into a gene list for identification of pathway modules. As a first step in the analysis, an input group of altered genes is mapped onto a literature-curated interaction network; several network sources are possible including Pathway Commons using the paxtoolsr package. In NetBox, candidate linker genes are defined as genes that they themselves do not have alterations but are direct neighbors of altered genes. To do this, in the second step, a hypergeometric test is used to determine the probability that a given candidate linker node has x or more interactions with nodes in the input gene list, *Pr*(*X≥x*), where x is the observed number of interactions between a candidate linker node to altered genes in the input list. This probability is taken as the p-value; significant p-values are suggestive that candidate linker genes are important to underlying biological processes along with the input genes. P-values of each candidate linker gene is corrected by the Benjamini-Hochberg method. Linker genes with an adjusted p-value equal to or less than 0.05 are counted as significantly connected linker genes. Third, a community detection algorithm (e.g., the Girvan–Newman edge betweenness algorithm) is applied to a new network that connects altered genes and significant connected linker genes from a literature-curated interaction network. NetBoxR offers edge-betweenness and leading eigenvector (for networks with large numbers of nodes) algorithm to identify network modules. Modules identified by NetBoxR can then be passed to enrichment packages, such as the ClusterProfiler package, to assign association of genes in the identified network modules to either canonical pathways or characterize the modules in terms of specific Gene Ontology (GO) terms (i.e., biological processes, molecular function, or cellular compartment).

### Assessment of statistical significance

Two statistical tests are performed on the identified network modules to assess significance. These tests were conducted in a similar manner as for the original Netbox algorithm (Cerami et al., 2010). To assess the level of global connectivity, an empirical p-value is calculated by determining the number of times the size of the largest network module identified from the same number of randomly selected genes equals or exceeds the size of the largest network module from the list of altered genes in the data set. Next, a network modularity score is calculated (Newman and Girvan, 2004). This score represents the strength (or quality) of the division of a network into various modules and is defined as the edge fraction that is within given modules minus an expected fraction by randomly distributing the edges. To assess the statistical significance of the network modularity observed in the resulting network, we used a local rewiring algorithm where random networks are generated that maintain the same size and all genes maintain the same degree, but the choice of interaction partners is random. For each of these random networks, we calculate the network modularity score and calculate the average and standard deviation for a set of random networks. The observed modularity score is then converted into a z-score (and reported as a p-value) to measure the deviation of the observed network modularity from that of the random null model.

### Implementation details and integration in the R ecosystem

Beyond the base functions, NetBoxR includes several additions to simplify and expand its use. These include 1) instructions for retrieving and processing genomic alterations such as SNV mutations and copy number alterations from data repositories such as cBioPortal (Cerami et al., 2012; Gao et al., 2013) or Genomic Data Commons (GDC) (Jensen et al., 2017), 2) ability to retrieve pathway data via the paxtoolsr package as available in ~20 data source aggregation Pathway Commons, (Luna et al., 2016) or STRING (Snel et al., 2000), 3) functionality to switch between the discovery of modules using various algorithms for detection of network communities from the igraph package (Csardi and Nepusz, 2005) and 4) guidance on annotating NetBoxR-derived modules using the ClusterProfiler package (Yu et al., 2012).

NetBoxR was implemented in R version 3.6.0 with igraph version 1.2.4.1 and should work in higher versions. The NetBoxR package can be installed through the BiocManager package manager for the Bioconductor package repository.

### Use Case

We tested the NetBoxR package in a cancer use case by using the list of altered genes from TCGA datasets and the prior-knowledge network data from the original Netbox paper (Cerami et al., 2010). The results reported here are comparable to those from Cerami et al. although the unadjusted p-values for linker genes are not exactly the same. This is because the unadjusted p-values of linker genes in the original NetBox report (Cerami et al., 2010) were calculated as the probability that a given candidate linker node has exactly X interactions Pr(X=x), where X is the observed number of interactions between a candidate linker node to altered genes in the input list; NetBoxR uses a hypergeometric test as described in the Base Functionality section. The final number of linker genes, using a significance cutoff of 0.05 after FDR correction, is the same between NetBoxR result and the original NetBox implementation. Using NetBoxR, we identify the RB1 and PIK3R1 modules, each with connected genes that are significantly altered as a set in the GBM data. Details for this example are provided in the vignette document in the NetBoxR package. Figure 1 demonstrates the use of the NetBoxR package to discover pathway modules from multiple genomic data for the example of the TCGA glioblastoma multiforme (GBM) study.

**Figure 1.**
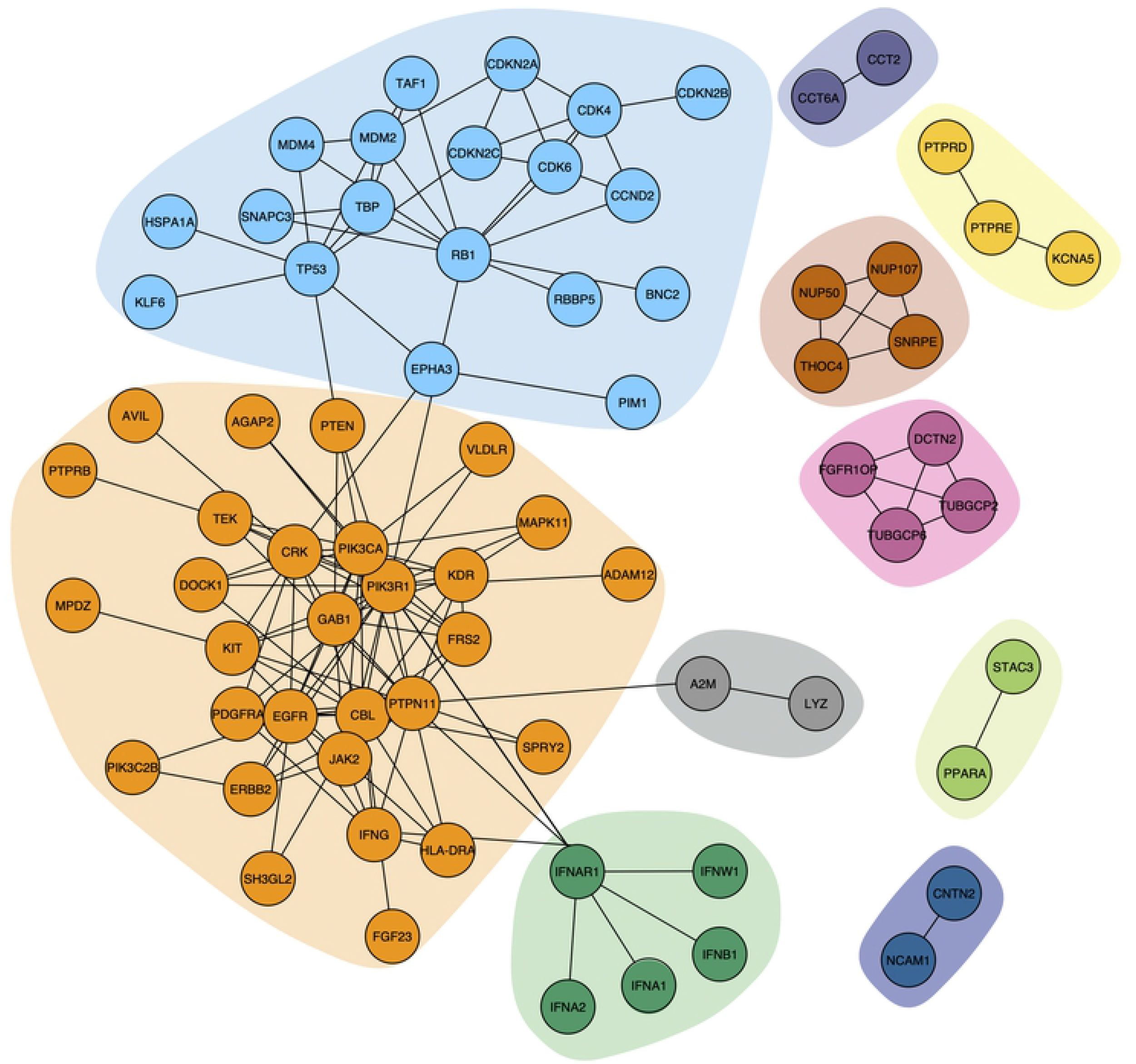
Glioblastoma multiforme (GBM) pathway modules identified by the NetBoxR package from cancer genomics alteration data without the use of pre-defined gene sets. The new NetBox algorithm implementation in R uses the igraph library to speed up module detection and visualization. Using mutations, and copy number alteration data from TCGA (Cerami et al., 2010), NetBoxR identified 10 pathway modules. The largest module (light orange background) contains genes related to the PIK3 pathway and the second largest module (light blue background) contains the genes related to the TP53 and cell cycle pathways.

## Conclusion

The NetBoxR R package facilitates data-driven network module discovery. Here we provide a use case that emphasizes analysis of cancer genomics data, but the methodology is applicable for other diseases or cell biological perturbation experiments with comparable large datasets covering genetic or molecular changes in genes or gene products. With the ease of installation in R bioinformatics environments, researchers can quickly use datasets of molecular measurements to identify pathway modules and form biological hypotheses on the functional role of cellular processes in cancer and other diseases, as well as in healthy tissues

## Availability of data and software

The NetBoxR software, source code, and tutorial are available in the Bioconductor repository: https://www.bioconductor.org/packages/release/bioc/html/netboxr.html.

Archived source at the time of publication: https://doi.org/doi:10.18129/B9.bioc.netboxr

License: LGPL-3

## Author contributions

EL, AL, and GD designed and wrote various parts of the Bioconductor package. EL, AL, and CS wrote the manuscript. CS developed the Netbox algorithm, in collaboration with Ethan Cerami, with further improvements by EL and AL.

## Acknowledgments

We would like to thank The Cancer Genome Atlas community of researchers for providing the genomic data used in the NetBoxR package vignette.

